# Genetic architecture of creativity and extensive genetic overlap with psychiatric disorders revealed from genome-wide association analyses of 241,736 individuals

**DOI:** 10.1101/2022.07.25.501322

**Authors:** Hyejin Kim, Yeeun Ahn, Joohyun Yoon, Kyeongmin Jung, Soyeon Kim, Injeong Shim, Tae Hwan Park, Hyunwoong Ko, Sang-Hyuk Jung, Jaeyoung Kim, Sanghyeon Park, Dong June Lee, Sunho Choi, Soojin Cha, Beomsu Kim, Min Young Cho, Hyunbin Cho, Dan Say Kim, Hong Kyu Ihm, Woong-Yang Park, Hasan Bakhshi, Kevin S O’Connell, Ole A Andreassen, Jonathan Flint, Kenneth S. Kendler, Woojae Myung, Hong-Hee Won

## Abstract

Creativity is heritable and exhibits familial aggregation with psychiatric disorders, but its genomic basis and genetic relationship with psychiatric disorders remain largely unknown. Here, we conducted a genome-wide association study (GWAS) using an expanded, machine learning-based definition of creativity in individuals of European ancestry from the UK Biobank (*n* = 241,736) and identified 25 creativity-associated loci. Extensive genetic overlap with psychiatric disorders, including schizophrenia, major depression, bipolar I disorder, attention deficit/hyperactivity disorder, and anorexia nervosa, was demonstrated by the genetic correlation, polygenic risk score, and MiXeR analyses. The condFDR and conjFDR analyses identified additional loci for creativity and psychiatric disorders, as well as shared genetic loci between creativity and psychiatric disorders. This GWAS showed significant correlations with GWASs using traditional definitions of creativity and GWASs adjusted for educational attainment. Our findings contribute to the understanding of the genetic architecture of creativity and reveal its polygenic relationships with psychiatric disorders.

## Introduction

Creativity is a multi-dimensional construct that encompasses aspects of cognitive processes, personality traits, and environmental factors^1, 2^. Creativity can be generally described as having novel ideas or alternatives to problems, exhibiting cognitive flexibility, and possessing the ability to synthesize as well as organize information. It is important to investigate creativity, as it is a significant trait that impacts individuals, businesses, and society. Artistic and literary professions, which were traditionally considered to be highly creative, have often been associated with psychopathology, which adds to the significance of creativity as a clinically relevant trait^3^.

The association between creativity and psychopathology has been theorized from ancient times, prompted by the anecdotal reports of artistic individuals exhibiting psychiatric symptoms^4, 5^. Reports of creative geniuses, such as Wolfgang Amadeus Mozart, Vincent van Gogh, and Edgar Allan Poe, exhibiting symptoms that seem to align with the diagnostic criteria of psychiatric illnesses has garnered attention from the public as well as academia. Since then, numerous researchers have attempted to provide empirical evidence of such association. Previous studies have utilized various methods to evaluate this link^6^, including assessing the rates of psychiatric disorders in individuals with creative occupations and their relatives, determining the likelihood of holding creative professions in psychiatric patients and their relatives, measuring creativity in relatives of psychiatric patients using interview-based scales^7^, and using polygenic risk scores (PRSs) of psychiatric illnesses to predict creativity^8^. The association between creativity and psychopathology has also been supported and summarized in large empirical reviews, with evidence for the co-segregation and polygenic relationship of creativity with mental illnesses^6, 9, 10^.

Nevertheless, there is a paucity of literature elucidating the genomic basis of creativity and its genetic relationship with psychiatric disorders. Although previous studies have shown that heritability of creativity is moderate to high^3^, the genetic architecture of creativity has not been revealed yet. Li *et al.*^11^ have examined the genetic architecture of creativity through a genome-wide association study (GWAS) but were not able to find any genome-wide significant loci due to a small sample size of 4,664 individuals. In addition, Li *et al.*^11^ did not perform single-nucleotide polymorphism (SNP)-based heritability and pathway analysis, nor analyses of genetic correlations and polygenic overlaps between creativity and psychiatric disorders.

In the present study, we conducted a GWAS of creativity using a large number of individuals to clarify the genetic architecture of creativity and its polygenic relationship with mental illnesses. To utilize the pre-existing information from the UK Biobank (UKB), we adapted a machine learning (ML) method developed by Bahkshi *et al*.^12^ to assign a probability of creativity to each occupation. The aims of this study are as follows: 1) to identify creativity-associated genetic variants and molecular mechanisms using the UKB data via GWAS and post-GWAS analyses; 2) to investigate the relationships between creativity and various traits, including psychiatric disorders, by using linkage disequilibrium score regression (LDSC), PRS analyses, and bivariate causal mixture modeling (MiXeR); 3) to explore the shared genetic basis of creativity and psychiatric disorders using the conditional and conjunctional false discovery rate (cond/conjFDR) approach; and 4) to validate our initial GWAS, which utilizes a novel definition of creativity, in comparison to those using traditional definitions of creativity and/or controlling for genetic effects of educational attainment.

## Results

### Creativity phenotyping

Various methods have been developed to measure the creativity of individuals^1^, one of them being the use of the individual’s occupation to define them as creative or not. Such method has been utilized in previous epidemiological^4, 6^ and genetic studies^8^. Therefore, we also sought to use occupation to assess creativity. To estimate creativity using pre-existing data from the UKB, we utilized the creativity probability dataset that was obtained via an ML-based method described by Bakhshi *et al.*^12^. In brief, Bakhshi *et al.*^12^ initially labeled 59 occupations as ‘creative’ and 61 occupations as ‘non-creative’, guided by the list of creative occupations specified by the Department of Digital, Culture, Media and Sport of the UK and detailed job descriptions of US Standard Occupational Classification (SOC) 2010 occupations provided by the O*Net database (https://www.onetcenter.org). Then, Bakhshi *et al.*^12^ developed a probabilistic classification algorithm using this training set and predicted the probability of creativity of all occupations in the US SOC 2010. After that, by matching US SOC 2010 codes with UK SOC 2010 codes, Bakhshi *et al.*^12^ predicted the creative probability of each UK SOC 2010 occupation. This ML-based method showed robust sensitivity and specificity (area under the receiver-operating characteristic curve range = [0.881, 0.958]) in various fields of research^13–15^.

A total of 241,736 individuals of European ancestry who answered the baseline occupation question in the UKB were included in our analysis (**Supplementary Table 1**). We matched the creative probability of each occupation in the UK SOC 2010 to occupations in the UK SOC 2000, according to the guidelines from the UK Office for National Statistics^16^. Using one-to-one and many-to-one matching approaches, we estimated the creative probability for each of the 351 occupation categories in the UKB and used it as the phenotype in this study (ranging from 0 to 1). Two UK SOC 2000 codes (1171, officers in armed forces; 3311, non-commissioned officers and other ranks) were excluded due to the lack of corresponding data in the UK SOC 2010. The creative probability of each occupation in the UKB is shown in **Supplementary Table 2**. The distribution of the probabilities in nine different occupational categories is presented in **Supplementary Fig. 1**.

In most previous studies using occupation to define creativity, rather than defining creativity as a continuous variable as we did for our main outcome, researchers utilized a dichotomous system of defining individuals who hold traditional artistic or scientific professions as creative and others as non-creative. We sought to evaluate whether our GWAS results using the ML-based method would mirror the GWAS results of the traditionally defined creativity. Thus, two more dichotomous variables were added to the participants’ data: traditionally creative occupations (narrowly defined artistic professions) and broadly defined artistic or scientific professions based on previous studies^6^. These creative classifications are also shown in **Supplementary Table 2**. We additionally adjusted for education years to evaluate the effect of educational attainment on creativity in our main model.

### Genetic architecture of creativity

#### Genome-wide significant association signals for creativity

We performed a GWAS for creativity of 241,736 European participants in the UKB (**Fig. 1**). A total of 25 lead SNPs at a genome-wide significant level (*P* < 5 × 10^-8^) were identified via linkage disequilibrium (LD) clumping (*r*^2^ < 0.2) with the 1000 Genomes Project European reference panel (hg19; **Table 1**). Regional plots of significant loci are presented in **Supplementary Fig. 2**. The quantile-quantile (Q-Q) plot of the GWAS demonstrates genomic inflation (λ = 1.30; **Supplementary Fig. 3**), which is attributable to their polygenicity (LDSC intercept = 1.077; *s.e.* = 0.009).

**Fig 1.**
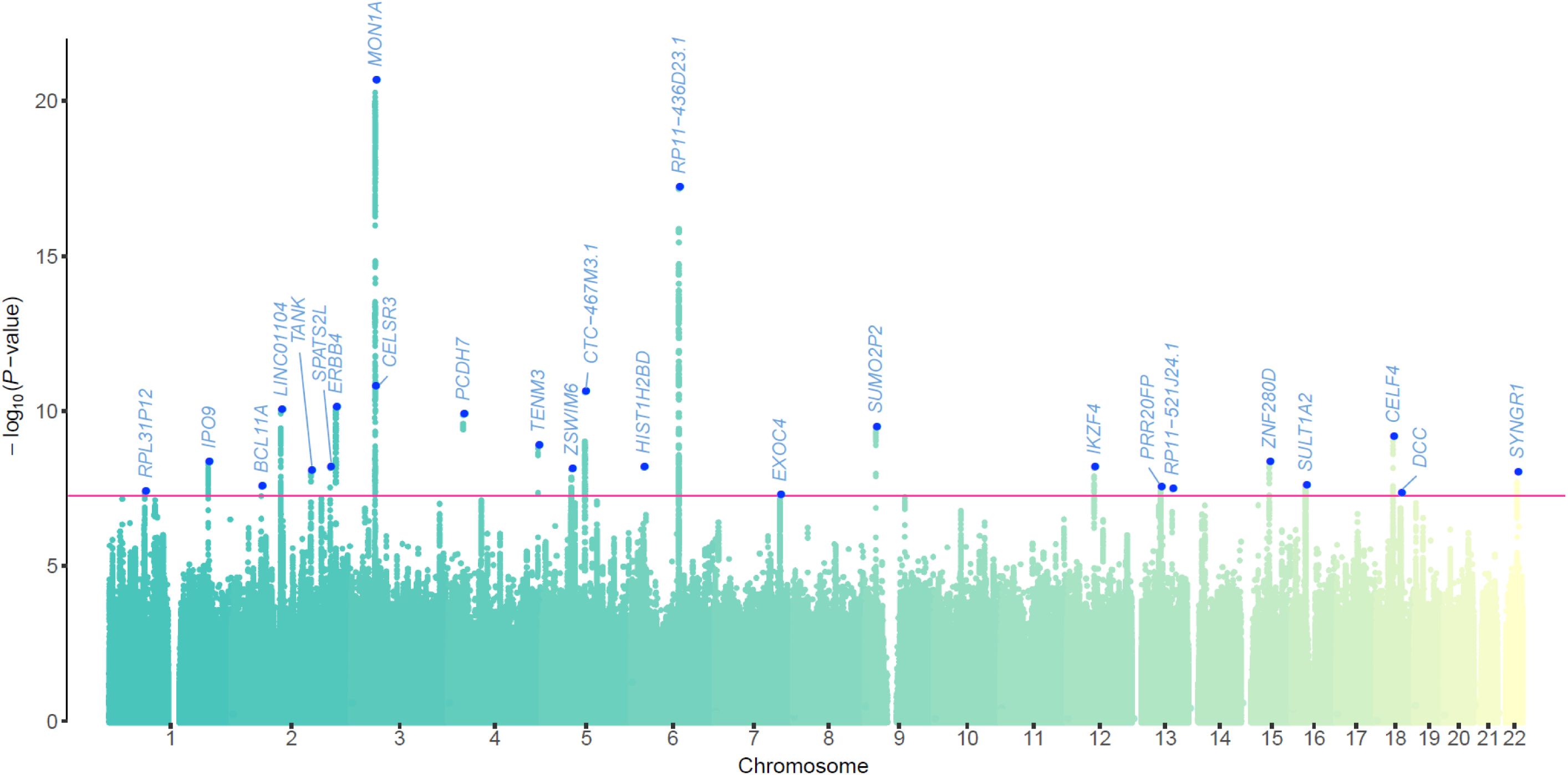
Manhattan plot for GWAS of creativity. The *x*-axis shows genomic positions and *y*-axis shows statistical significance as −log_10_ (*P*) values. The threshold for significance, which accounts for multiple tests, is indicated by the red horizontal line (*P =* 5 × 10^−8^). The blue dot indicates the nearest mapped gene from the lead SNPs.

**Table 1.**
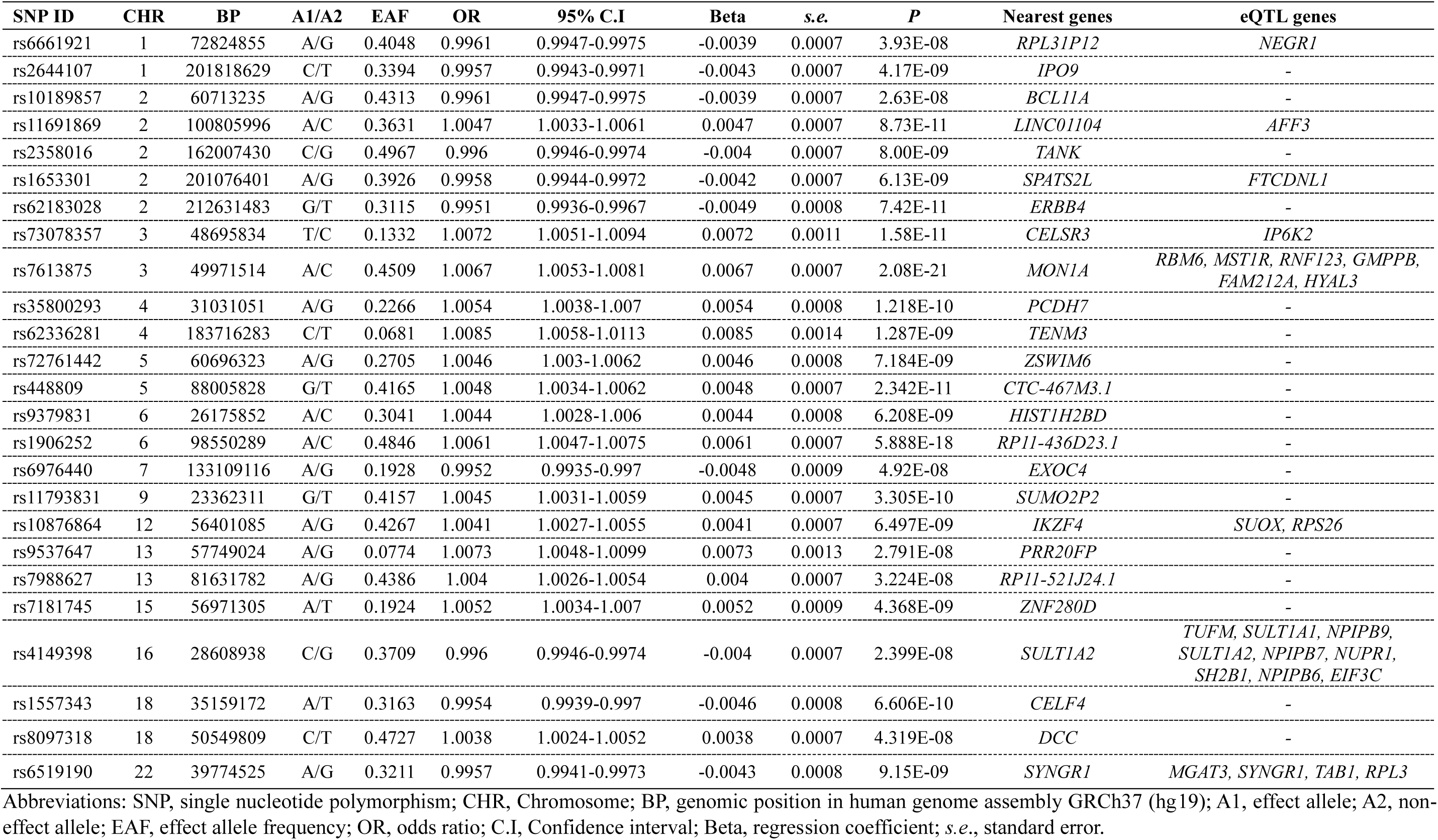
Summary of the lead SNPs in the 25 loci associated with creativity.

#### Functional annotation and biological pathways

We performed functional annotation using the Genotype-Tissue Expression (GTEx) database to explore relevant genes for the identified variants. Through expression quantitative trait loci (eQTL) analysis, a total of 25 cis-eQTL genes were mapped to the lead SNPs in the 13 brain tissue types (**Supplementary Table 3**). Based on the mapped eQTL genes, eight lead SNPs were identified, including rs6661921, rs11691869, rs1653301, rs7613875, rs73078357, rs10876864, rs4149398, and rs6519190 (**Table 1**). We also conducted biological pathway analysis, and found that a total of 678 biological pathways related to creativity were enriched (FDR-corrected *P* < 0.05; **Supplementary Table 4**). Pathways of forebrain neuron differentiation, forebrain generation of neurons, and guanosine diphosphate binding were significantly enriched.

#### SNP heritability and partitioned heritability analysis

We estimated SNP-based and partitioned heritability to explore the effect of total SNPs on creativity and to evaluate the enrichment of 53 genomic annotations. The total SNP heritability of creativity was estimated to be 8.62% (*s.e.* = 0.4%). Among the 53 annotations, only that of conserved genomic regions defined in the study by Lindblad-Toh *et al*.^17^ was significant for creativity at an FDR < 0.05 (**Fig. 2a** and **Supplementary Table 5**). The proportion of SNP heritability for the conserved regions was 2.6% and the estimated enrichment value was >15 (coefficient *P* = 4.56 × 10^−9^).

**Fig 2.**
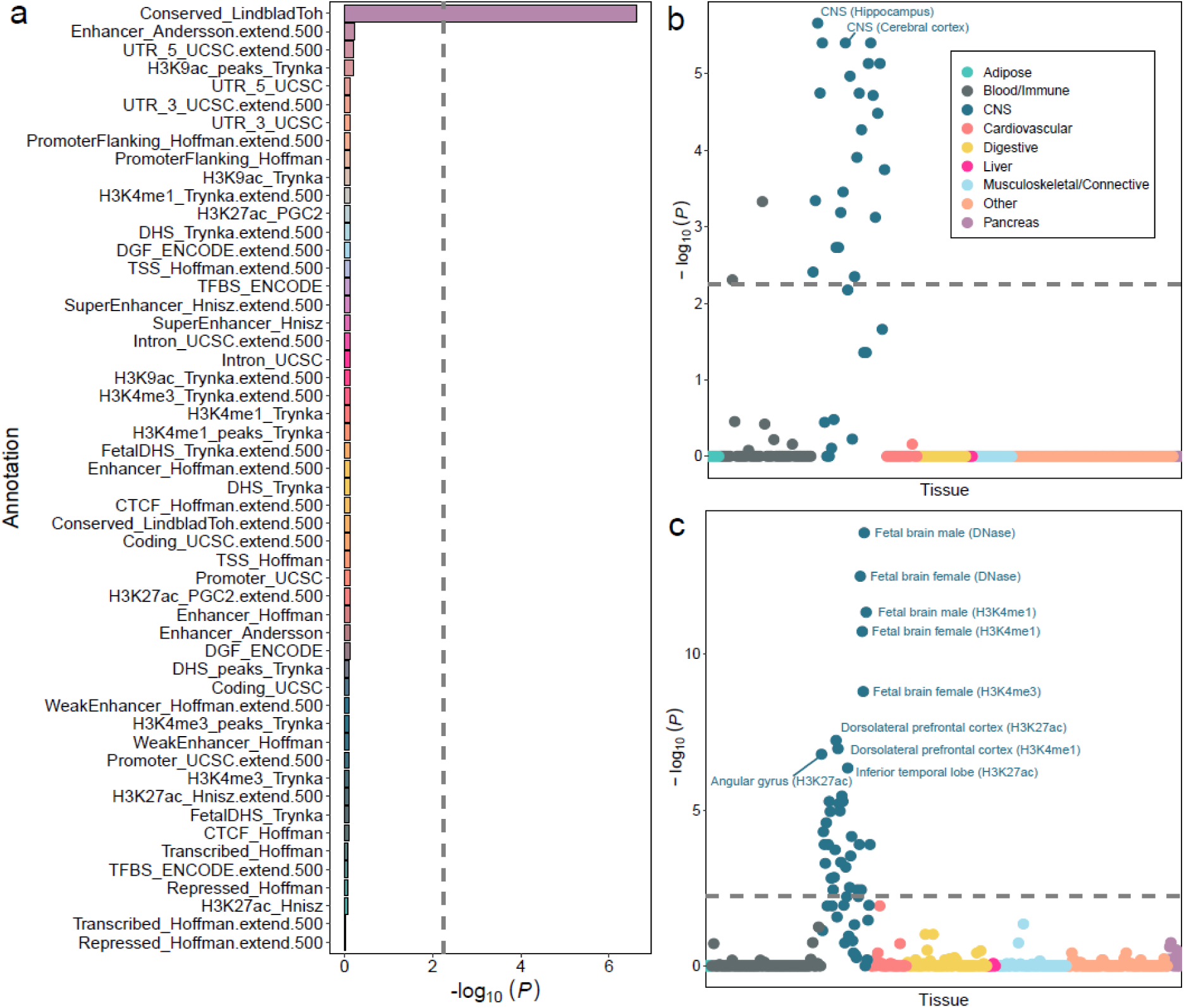
Partitioned heritability analyses using LDSC. **a.** Enrichment estimates for 53 functional annotations. Annotations are ordered by their *P* values. The dashed line indicates the significance at an FDR-corrected *P* < 0.05. **b.** Results of multiple-tissue analysis using gene expression data. Each circle represents a tissue or cell type from either the GTEx dataset or Franke lab dataset. The dashed line indicates the cutoff of FDR, which is <5% at a −log_10_ (*P*) = 2.22. **c.** Results of multiple-tissue analysis using chromatin data. Each circle represents a peak for DNase I hypersensitivity (DHS) or histone marks in a tissue or cell type. The dashed line indicates the cutoff of FDR, which is <5% at a −log_10_ (*P*) = 2.37.

In the LDSC applied to specifically expressed gene (LDSC-SEG) analysis for creativity across multiple tissues, we found that the GWAS signals were strongly enriched in the hippocampus and cerebral cortex in the central nervous system (CNS) (**Fig. 2b** and **Supplementary Table 6** for all other significant tissues). In the multi-tissue chromatin results, we observed strong enrichment of dorsolateral prefrontal cortex, angular gyrus, and inferior temporal lobe in the CNS (**Fig. 2c** and **Supplementary Table 7** for all other significant tissues). The CD4^+^ T cell gene set (T.4int8+.Th; genes expressed in T-helper cells with CD4^+^, intermediate CD8, and high T-cell receptors) and neurons were also strongly enriched among immune cell types and CNS tissues, respectively, at an FDR < 0.05 (**Supplementary Tables 8** and **9**).

### Genetic relationships between creativity and other traits

#### Genetic correlation of creativity with other traits

We examined the genetic correlations between creativity and health-related traits (**Fig. 3** and **Supplementary Table 10**). A significant positive correlation was observed between creativity and traits of high-density lipoprotein (HDL) cholesterol (*r*_g_ = 0.25, *P_FDR* = 6.31 × 10^−27^), testosterone (*r*_g_ = 0.10, *P_FDR* = 3.37 × 10^−5^), never smoking (*r*_g_ = 0.32, *P_FDR* = 1.18 × 10^−42^), sleep duration (*r*_g_ = 0.17, *P_FDR* = 2.47 × 10^−10^), eveningness chronotype (*r*_g_ = 0.12, *P_FDR* = 0.0004), miscarriage (*r*_g_ = 0.35, *P_FDR* = 0.0233), worry too long after embarrassment (*r*_g_ = 0.18, *P_FDR* = 4.56 × 10^−9^), fluid intelligence test-related score and sub-types (*r*_g_ range = [0.50, 0.78], *P_FDR* range = [1.98 × 10^−223^, 1.70 × 10^−16^]), Parkinson’s disease (*r*_g_ = 0.14, *P_FDR* = 0.0003), anorexia nervosa (AN, *r*_g_ = 0.22. *P_FDR* = 2.59 × 10^−6^), and bipolar I disorder (BD I, *r*_g_ = 0.12, *P_FDR* = 0.0003). A significant negative correlation was observed with vitamin D (*r*_g_ = −0.11, *P_FDR* = 8.46 × 10^−7^), obesity (*r*_g_ = −0.44, *P_FDR* = 2.63 × 10^−34^), time spent watching television (*r*_g_ = −0.67, *P_FDR* = 194 × 10^−228^), duration of walks (*r*_g_ = −0.70, *P_FDR* = 4.86 × 10^−131^), sleep disorders (*r*_g_ = −0.19 *P_FDR* = 0.0003), urinary tract infection (*r*_g_ = −0.57, *P_FDR* = 3.47 × 10^−10^), loneliness (*r*_g_ = −0.30, *P_FDR* = 7.22 × 10^−26^), fed-up feelings (*r*_g_ = −0.43, *P_FDR* = 3.03 × 10^−62^), ever unenthusiastic/disinterested for a whole week (*r*_g_ = −0.19, *P_FDR* = 7.29 × 10^−5^), anxiety disorders (*r*_g_ = −0.37, *P_FDR* = 9.19 × 10^−11^), financial situation satisfaction (*r*_g_ = −0.44, *P_FDR* = 5.92 × 10^−24^), Alzheimer’s disease (*r*_g_ = −0.23, *P_FDR* = 0.0068), major depression (MD, *r*_g_ = −0.20, *P_FDR* = 1.17 × 10^−13^), and attention deficit/hyperactivity disorder (ADHD, *r*_g_ = −0.48, *P_FDR* = 7.96 × 10^−40^). These results suggest that various health-related traits share genetic bases with creativity.

**Fig 3.**
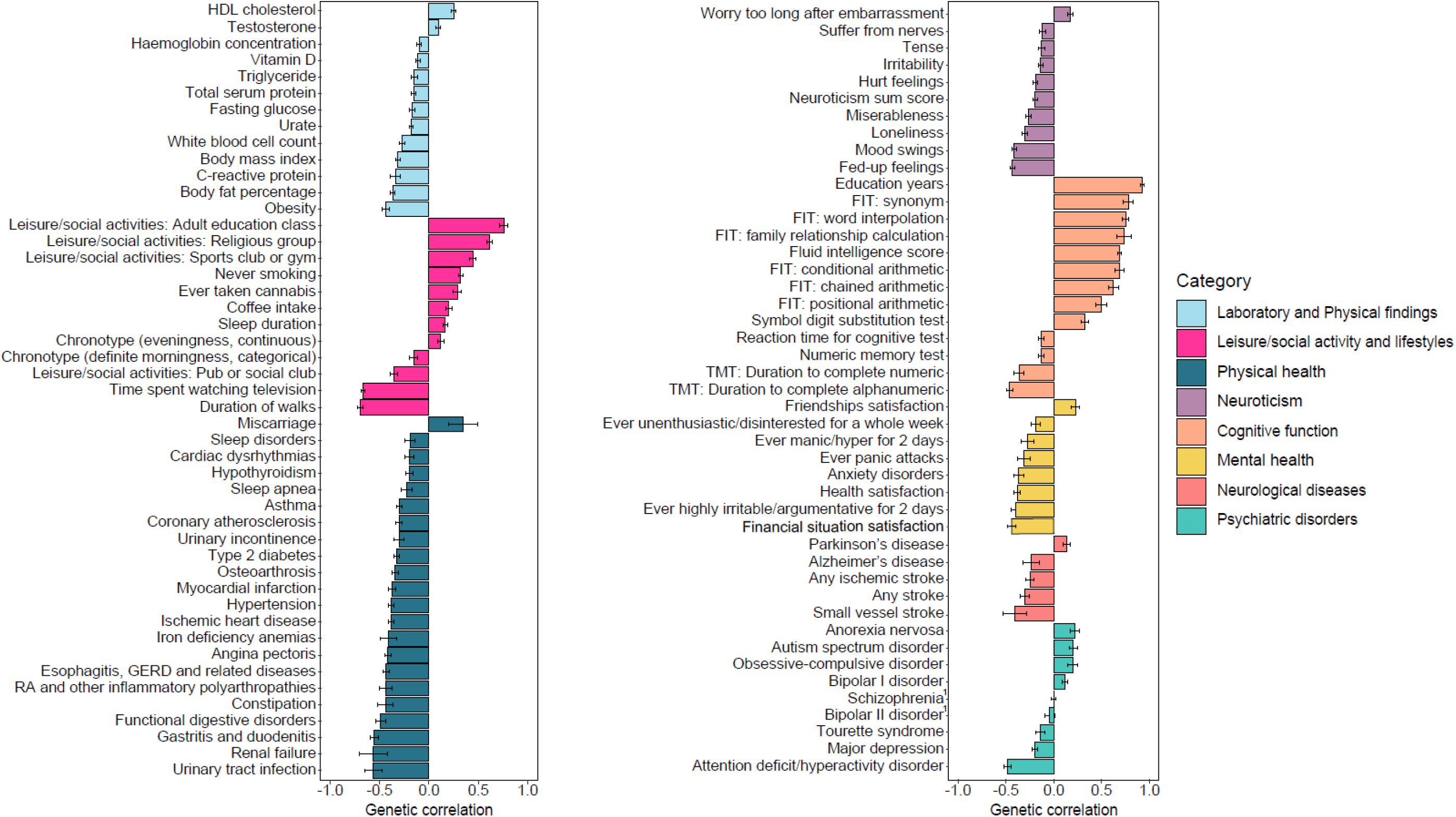
Genetic correlation estimates between creativity and other phenotypes using LDSC. This figure includes significant genetic correlations where FDR values are <5% (see **Supplementary Table 10** for all results). Two traits (schizophrenia and bipolar II disorder) were not significant, but are included in this figure as an exception for psychiatric disorders. Abbreviations: HDL, high-density lipoprotein; GERD, gastroesophageal reflux disease; FIT, fluid intelligence test; TMT, trail making test.

We also conducted LDSC to estimate the genetic correlation between creativity and neuroimaging traits, such as brain volume and diffusion tensor imaging (DTI) measures (**Supplementary Table 11**). After FDR correction, we observed that only the total brain volume demonstrated a significant positive association with creativity (*r*_g_ = 0.19).

#### Polygenic risk scores for psychiatric disorders and associations with creativity

With the PRS analyses, we investigated associations of creativity with psychiatric disorders using the summary statistics of GWAS results (**Supplementary Table 12** and **Fig. 4**). PRSs for nine psychiatric disorders were significantly associated with creativity. Of them, PRS of ADHD showed the largest R^2^, explaining approximately 0.25% of the variance of creativity. We also observed positive relationships between creativity and the PRSs for BD I (coefficient = 96.43, *s.e.* = 13.30, *P* = 4.22 × 10^−13^), autism spectrum disorder (ASD; coefficient = 50.62, *s.e.* = 6.81, *P* = 1.07 × 10^−13^), AN (coefficient = 177.11, *s.e.* = 18.77, *P* = 3.93 × 10^−21^), and obsessive compulsive disorder (OCD; coefficient = 47.75, *s.e.* = 5.83, *P* = 2.51 × 10^−16^) and negative relationships between creativity and the PRSs for SCZ (coefficient = –65.09, *s.e.* = 19.34, *P* = 0.001), MD (coefficient = –687.83, *s.e.* = 38.44, *P* = 1.42 × 10^−71^), bipolar II disorder (BD II; coefficient = –0.04, *s.e.* = 0.02, *P* = 0.037), ADHD (coefficient = –361.94, *s.e.* = 14.74, *P* = 5.11 × 10^−133^), and Tourette syndrome (TS; coefficient = –13.95, *s.e.* = 3.85, *P* = 0.0003).

**Fig 4.**
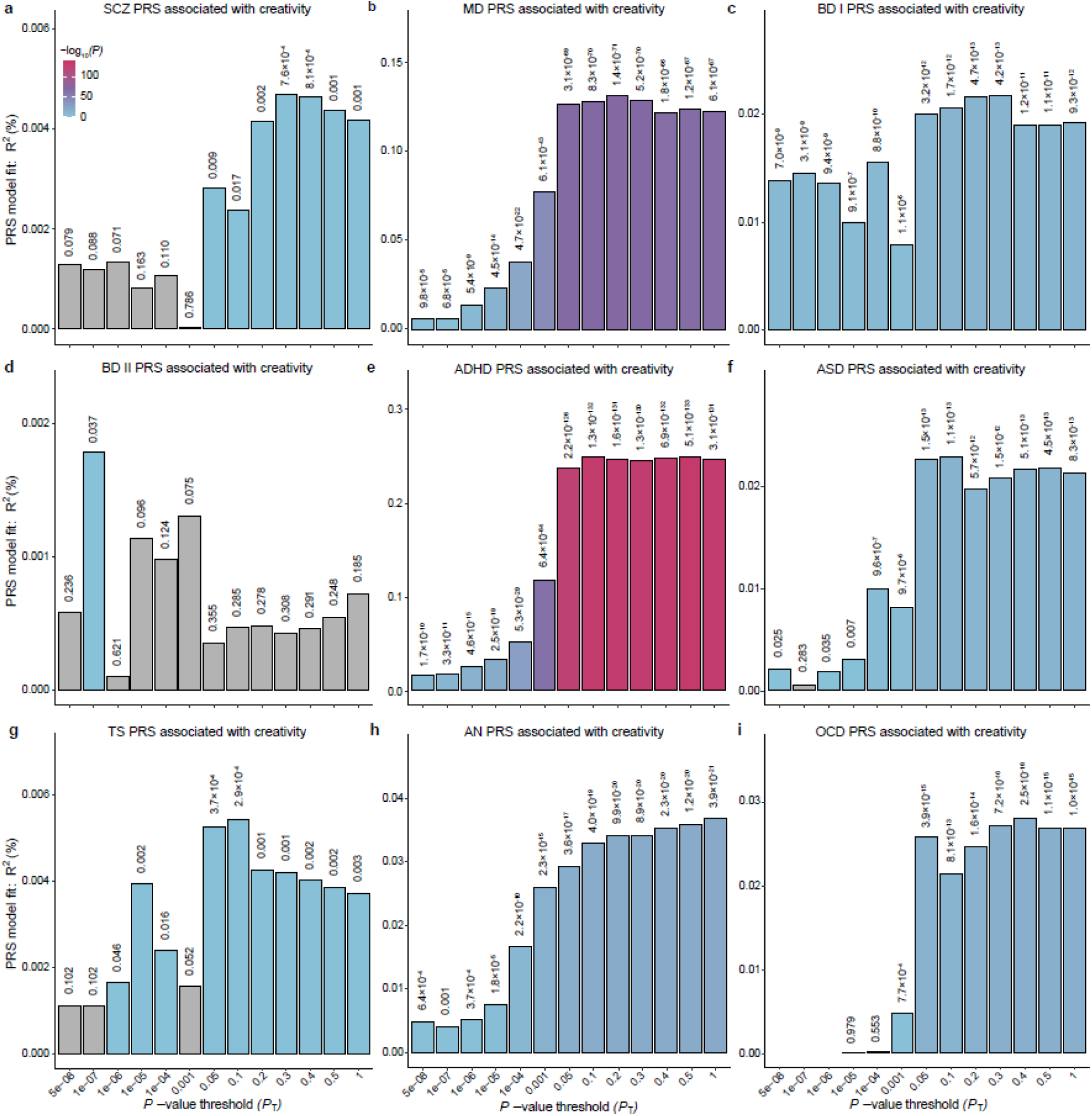
Polygenic risk scores for psychiatric disorders associated with creativity (ML-based creative probability). Polygenic risk scores for psychiatric disorders associated with creativity. Nagelkerke’s pseudo-*R*^2^ (*y*-axis) is shown for scores derived using 13 thresholds ranging from 5 × 10^−8^ to 1 (*x*-axis). The significance increases from blue to red, and grey represents non-significance. The *P*-value is specified above each bar. **a.** SCZ PRS associated with creativity. **b.** MD PRS associated with creativity. **c.** BD I PRS associated with creativity. **d.** BD II PRS associated with creativity. **e.** ADHD PRS associated with creativity. **f.** ASD PRS associated with creativity. **g.** TS PRS associated with creativity. **h.** AN PRS associated with creativity. **i.** OCD PRS associated with creativity. Abbreviations: PRS, polygenic risk score.

#### Polygenic overlap between creativity and psychiatric disorders

We estimated the polygenic overlap between creativity and psychiatric disorders using MiXeR (**Supplementary Fig. 4** and **Supplementary Tables 13** and **14**). Based on the positive value of the Akaike Information Criterion (AIC), the results of SCZ, MD, BD I, ADHD, and AN were determined to be reliable (**Fig. 5a**). MD showed the most genetic overlap with creativity (Dice coefficient; DC = 0.94), sharing approximately 11,100 out of 12,600 causal variants. This indicates that most of the SNPs affecting creativity also affect MD. Similarly, BD I (DC = 0.74), ADHD (DC = 0.67), and AN (DC = 0.78) shared approximately 59% (7,000 out of 11,800), 50% (5,800 out of 11,500), and 64% (7,400 out of 11,600) of SNPs with creativity, respectively, demonstrating considerable polygenic overlap between creativity and the three disorders. Interestingly, it was found that SCZ (DC = 0.89) also shared a substantial portion of SNPs with creativity (approximately 9,300 out of 11,700) despite the weak genetic correlation found in the LDSC analysis.

**Fig 5.**
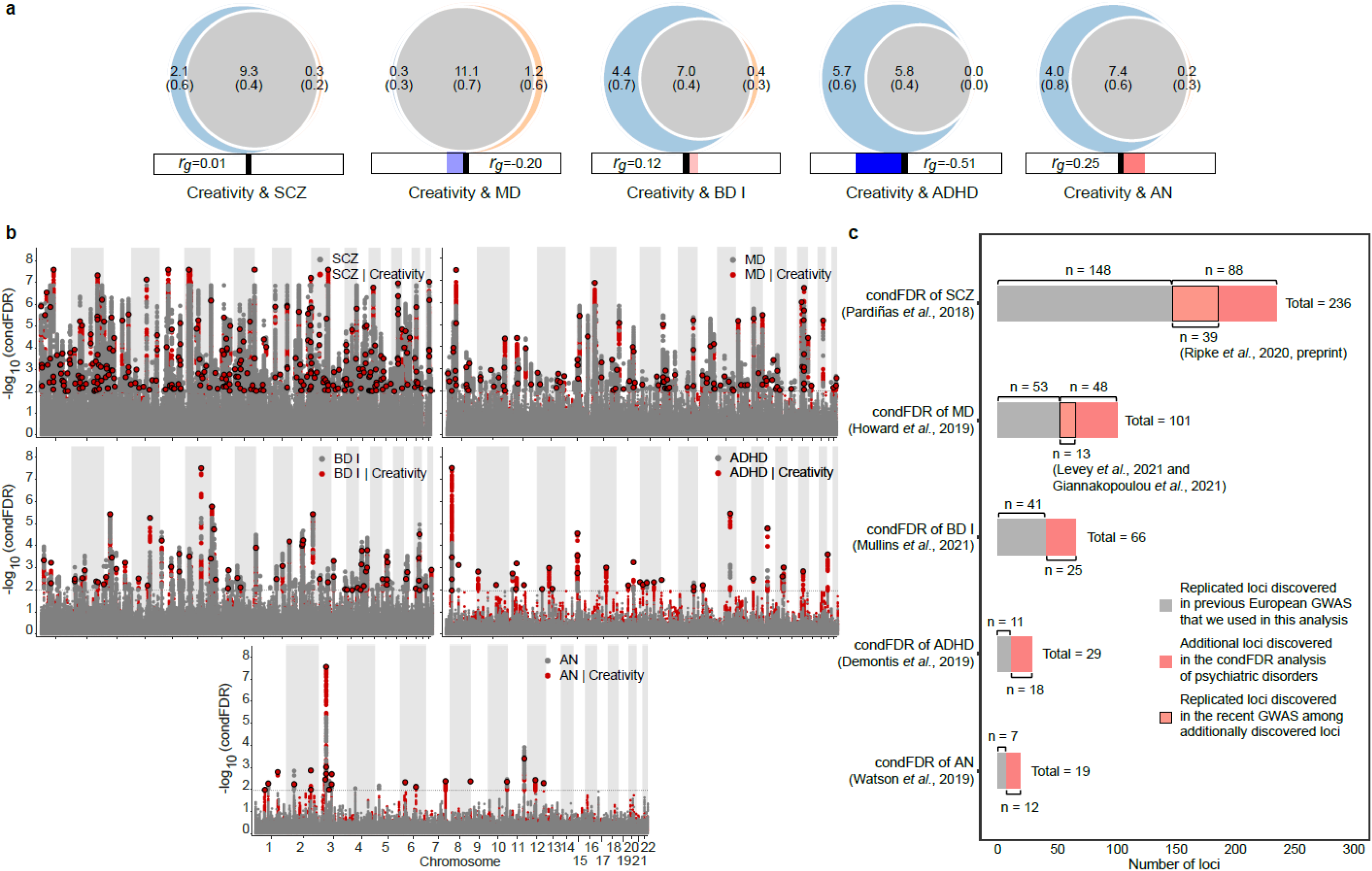
Polygenic overlap between creativity and psychiatric disorders (schizophrenia, major depression, bipolar I disorder, attention deficit/hyperactivity disorder, and anorexia nervosa). **a.** Venn diagrams depicting the estimated number of trait-influencing variants shared (grey) between creativity (left circle) and psychiatric disorders (right circle; schizophrenia, major depression, bipolar I disorder, attention deficit/hyperactivity disorder, and anorexia nervosa) and unique (colors) to either of them (see **Supplementary Fig. 4** for all results). The number of trait-influencing variants in thousands is shown, with the standard error in thousands given in parentheses. The estimated genetic correlation for each pair is indicated below the corresponding Venn diagram, with an accompanying directional scale (blue shades for negative scale and red shades for positive scale). **b.** Conditional Manhattan plots of −log_10_ scale of condFDR values for psychiatric disorders alone and psychiatric disorders given creativity. SNPs with −log_10_ (condFDR) > 2 (*i.e.*, FDR < 0.01) are indicated by large circles. A black line around the large circle indicates the most significant SNP in each linkage disequilibrium block. **c.** Significant condFDR variants stratified and compared with previously reported variants. A total of 88 variants for SCZ, 48 variants for MD, 25 variants for BD I, 18 variants for ADHD, and 12 variants for AN were identified in addition to those obtained from the GWAS summary statistics used in this study (see **Supplementary Tables 15, 16** and **17**). Among the 88 SCZ variants, 39 were replicated in the latest GWAS by Ripke *et al*.^23^. Among the 48 MD variants, 13 were replicated in two GWAS results by Levey *et al*.^24^ and Giannakopoulou *et al*.^25^.

### Conditional and conjunctional FDR and functional annotation

#### Conditional FDR for psychiatric disorders and creativity

For the condFDR analyses of SCZ, MD, BD I, ADHD, and AN conditional on creativity, we used a conditional Manhattan plot to illustrate the localization of the genetic variants (**Fig. 5b**). We identified 236 SCZ-related genomic loci conditional on its associations with creativity. Of these loci, 88 were additionally identified for SCZ compared with a previous study^18^. In the condFDR analysis of MD, we identified 101 MD-associated loci, of which 48 were additional^19^. In the condFDR analysis of BD I, we identified 66 BD I-related loci, of which 25 were additional compared to those identified in a previous GWAS^20^. The condFDR analysis of ADHD and AN identified 29 and 19 loci, respectively, of which 18 and 12 loci were additional in comparison to previous GWAS results^21, 22^ (**Fig. 5c** and **Supplementary Tables 15–19**). Furthermore, we examined whether the additional 88 loci associated with SCZ were replicated in the latest cross-ancestry GWAS analysis of Ripke *et al*.^23^ Of the 88 loci, 39 satisfied the genome-wide significant level in the GWAS analysis of Ripke *et al*. and the other 49 were not reported. We also examined whether the additional 48 loci associated with MD were replicated in the two recent GWAS analyses of Levey *et al*.^24^ and Giannakopoulou *et al*.^25^. Among the 48 loci, 13 were genome-wide significant in the two studies and the other 35 were not reported.

For these additionally identified loci of each psychiatric disorder in our study (88 loci for SCZ, 48 loci for MD, 25 loci for BD I, 18 loci for ADHD, and 12 loci for AN), 100, 27, 29, 14, and 5 genes were mapped to the loci by eQTL analysis, respectively (**Supplementary Tables 20–24**). The 100 mapped genes for SCZ were enriched in two specific tissue types, namely the cerebellar hemispheres and cerebellum (**Supplementary Fig. 5.1**). The 29 mapped genes associated with BD I were enriched in four brain tissue types including the cortex and cerebellum (**Supplementary Fig. 5.3**). However, for the MD, ADHD, and AN mapped genes, there was no significant enrichment in brain tissue (**Supplementary Figs. 5.2, 5.4,** and **5.5**).

We also sought to identify additional loci associated with creativity beyond our initial GWAS using the condFDR analyses conditional on psychiatric disorders (**Supplementary Fig. 6** and **Supplementary Table 25**). We discovered 42, 48, 43, 40, and 42 creativity-associated genomic loci using SCZ, MD, BD I, ADHD, and AN as the associated phenotypes, respectively. Of these loci, 17, 24, 18, 18, and 15 loci, respectively, were not detected by our initial GWAS. Overall, 69 additional loci for creativity were identified.

For the additional loci of creativity given psychiatric disorders (**Supplementary Table 25**), eQTL mapping was performed (**Supplementary Table 26**). We identified 56, 44, 61, 42, and 11 genes mapped to the creativity-associated additional loci using SCZ, MD, BD I, ADHD, and AN, respectively. The 56 mapped genes given SCZ were significantly and differentially expressed in the brain cerebellum. The 61 mapped genes using BD I were enriched in two specific tissue types, including the cerebellum and cerebellar hemispheres. However, there were no enriched brain tissues associated with the mapped genes using MD, ADHD, and AN (**Supplementary Fig. 7**).

#### Conjunctional FDR between creativity and psychiatric disorders

For the conjFDR analyses between creativity and psychiatric disorders, we used a conjunctional Manhattan plot to present the distribution of significant genetic variants (**Supplementary Fig. 8**). We identified 50 shared genomic loci between creativity and SCZ, 80 loci between creativity and MD, 26 loci between creativity and BD I, 33 loci between creativity and ADHD, and 21 loci between creativity and AN (**Supplementary Table 27**).

For these jointly associated genomic loci between creativity and psychiatric disorders, 119 genes between creativity and SCZ, 100 genes between creativity and MD, 34 genes between creativity and BD I, 32 genes between creativity and ADHD, and 38 genes between creativity and AN were mapped using eQTL analysis (**Supplementary Table 28**). The 119 mapped genes for creativity and SCZ were enriched in three brain tissue types, including the cerebellum, putamen, and basal ganglia. Brain tissues were also enriched for the mapped genes between creativity and other psychiatric disorders, except for the genes mapped for AN (**Supplementary Fig. 9**).

#### Comparison with GWAS of dichotomous definitions of creativity and GWAS adjusted for educational attainment

For sensitivity analyses, we additionally performed two GWASs using dichotomous definitions of creativity: narrowly defined artistic professions and broadly defined artistic or scientific professions (**Supplementary Table 2** and **Supplementary Figs. 10** and **11**). The initial GWAS showed strong positive genetic correlations with the two GWASs of dichotomous creative phenotypes: narrowly defined artistic professions (*r*_g_ = 0.73) and broadly defined artistic or scientific professions (*r*_g_ = 0.90). In line with the results of genetic correlation, the effect size correlation of the significant SNPs between the initial GWAS and GWASs of traditionally defined creativity were high (*rho* range = [0.87, 0.94], *P* < 2.2 × 10^−16^; **Supplementary Figs. 13.1-13.3**). In the PRS analysis using the GWAS of narrowly defined artistic professions (**Supplementary Table 29**), the direction of most association signals with psychiatric disorders was consistent with those using the initial GWAS except for SCZ, based on the largest R^2^ and *P* value.

The initial GWAS for ML-based definition of creativity adjusted for education years is depicted in Manhattan and Q-Q plots (**Supplementary Figs. 12.1** and **12.2**), indicating a complete concordance with the initial GWAS (*r*_g_ = 1.00 and *rho* = 1.00, *P* < 2.2 × 10^−16^ in effect size correlation analysis). The two GWASs using dichotomous creative phenotypes were also additionally adjusted for education years (**Supplementary Figs. 12.3-12.6**) and showed a complete concordance of significant SNPs with the two GWASs before adjusting for education years (**Supplementary Figs. 13.4-13.5**), along with a perfect genetic correlation (*r*_g_ = 1.00).

## Discussion

In this study, we performed the largest GWAS investigating creativity to date in which we utilized the creativity probability dataset that was obtained via an ML-based method to measure the creativity of individuals from the UKB. We performed various genomic analyses to clarify the genetic basis of creativity and its relationship with psychiatric disorders. Our initial GWAS identified 25 lead variants associated with creativity in the UKB participants (**Fig. 1** and **Table 1**). The heritability of all SNPs was estimated to be 8.62%, indicating that the analyzed creativity phenotype had a significant genetic component. Through eQTL and LDSC-SEG analyses, we discovered that creativity was strongly associated with the CNS, specifically neurons (**Fig. 2b, 2c, Supplementary Tables 6** and **9**). Additionally, we revealed significant genetic correlations between creativity and various health-related traits as well as cognitive function, neurological diseases, and psychiatric traits and disorders using LDSC analysis (**Fig. 3**). Genetic associations between creativity and psychiatric disorders were supported by the results of PRS (**Fig. 4**) and MiXeR analyses (**Fig. 5a**), specifically emphasizing the polygenic overlap of creativity with SCZ, MD, BD I, ADHD, and AN. The condFDR and conjFDR analyses provided further insights into the genetic overlap between creativity and psychiatric disorders by identifying additional and shared SNPs (**Fig. 5b, 5c, Supplementary Tables 15-19** and **27**). Moreover, our initial GWAS showed strong positive genetic correlations with GWASs of narrowly and broadly defined creative professions, demonstrating that our novel definition of creativity (ML-based creative probability) can be considered as an expanded, validated measure of creativity that can provide more robust results. The GWASs adjusted for education years also showed a complete concordance with the GWASs before educational adjustment, confirming that educational attainment did not largely affect the GWAS results.

A recent GWAS conducted by Li *et al.*^11^ defined creativity based on a self-report questionnaire using 4,664 Han Chinese subjects. However, it is practically difficult to obtain creativity scores through questionnaires or evaluations in a large sample. Since occupations have been frequently used to define creativity^4–6, 8^ and we aimed to conduct our genomic analyses with the large, pre-existing data of the UKB, we utilized the creativity probability dataset obtained via an ML-method by Bakhshi *et al.*^12^ as an alternative way to define creativity. Using this method, we identified 25 lead variants that reached a genome-wide significance level (**Fig. 1, Table 1**). In addition, eight lead SNPs were identified as eQTLs associated with 25 genes in 13 brain tissue types: rs6661921, rs11691869, rs1653301, rs7613875, rs73078357, rs10876864, rs4149398, and rs6519190. Both rs73078357 and rs6519190 loci are located in the intron regions of *RBM6* and *SYNGR1*, respectively, which are associated with overall cognitive performance^26^ and SCZ^27^. *NEGR1*, which is located near the rs6661921 locus, is associated with MD and ADHD^28, 29^. *AFF3,* located near the rs11691869 locus, is a transcription activator that binds to double-stranded DNA, and has been previously associated with SCZ^27^ as well as intellectual disability^30^, and is predictive of general cognitive functioning^26^. *FTCDNL1*, positioned near the rs1653301 locus, is related to SCZ^31^. *IP6K2*, located near the rs7613875 locus, was reported to be involved in the physiology of SCZ^32^ and ADHD^33^. *RPS26* and *SULT1A2*, adjacent to the rs10876864 and rs4149398 loci, are associated with AN^34^ and ASD^35^, respectively. Several of these genes are also associated with risk-taking and psychiatric disorders, which is in accordance with previous studies^4, 6, 36^.

LDSC-SEG analysis, in addition to eQTL analysis that demonstrated the involvement of eQTL genes and brain tissues in creativity, revealed that brain tissues and neurons are significantly enriched for creativity. The enrichment analysis showed a broad involvement of brain tissues, including the hippocampus, cerebral cortex, and limbic system, compared to other types of tissues (**Fig. 2b, Fig. 2c**, and **Supplementary Table 6**), and exhibited enrichment within neurons rather than within oligodendrocytes and astrocytes (**Supplementary Table 9**). Conserved genomic regions defined by Lindblad-Toh *et al*.^17^, which were the only significant annotation among the 53 functional genomic annotations, contributed to approximately 2.57% of the total SNP heritability (8.62%, **Fig. 2a, Supplementary Table 5**), consistent with previous studies investigating SCZ, BD, AN, and educational attainment^37^. Our findings suggest that the brain is profoundly involved in the biological mechanisms of creativity, which aligns with previous studies on creativity and brain activity^38, 39^. Moreover, in accordance with findings of previous studies^40–42^, our genetic correlation analysis not only identified positive correlations between creativity and educational years, cognitive ability, as well as BD I (**Fig. 3** and **Supplementary Tables 10** and **11**), but also identified positive correlations between creativity and ASD, AN, and OCD as well as negative correlations with MD, ADHD, and TS. However, the LDSC analysis did not detect any significant correlations between creativity and SCZ or BD II.

The genetic correlations between creativity and psychiatric disorders were further supported by the results of PRS (**Fig. 4** and **Supplementary Table 12**) and MiXeR analyses (**Fig. 5a** and **Supplementary Tables 13** and **14**). The genetic relationships between creativity and psychiatric disorders based on the LDSC, PRS, MiXeR, condFDR, and conjFDR approaches are summarized in **Supplementary Table 30**. Based on the significant thresholds from PRS analyses, the coefficients of each PRS for psychiatric disorders generally showed the same trends with the results from the LDSC analyses, except for ASD (**Fig. 4**). The PRS for ADHD demonstrated the most significant association with creativity, explaining a maximum of 0.25% of the variance. Previous findings on the relationship between creativity and ADHD have been mixed^43^; however, our results indicate that ADHD may have the strongest, negative genetic influences on creativity among the nine psychiatric disorders analyzed. Consistent with a previous study^8^, our results showed that BD I PRS was positively associated with creativity, but not BD II PRS. This might be due to the differential effects of BD I and BD II symptoms on creativity^42^. While an association between creativity and the PRSs of SCZ and MD has previously been reported by Li *et al*.^11^, these associations were found to be mixed in our results; Li *et al*.^11^ showed a positive association of creativity with SCZ PRS as well as MD PRS while we identified these PRSs to be negatively associated with creativity when using the ML-based definition (**Supplementary Table 12**), but not when using narrowly defined artistic professions (**Supplementary Table 29**). Our speculation is that individuals who work in artistic occupations with higher creative probability (*i.e.*, those who are more creative) may have different potential risks for psychiatric disorders than those who work in scientific or other occupations with relatively lower creative probability (*i.e.*, those who are less creative). Low genetic correlation but extensive genetic overlap between creativity and SCZ may also reflect the existence of numerous genetic variants with different directional effects, complicating the shared genetic architecture between them. Furthermore, creativity measured via different methods could derive differential associations with psychiatric disorders, particularly SCZ. Nevertheless, further studies are necessary to clarify the nature of the associations between creativity and SCZ PRS and MD PRS. The MiXeR analysis indicated that SCZ, MD, BD I, ADHD, and AN showed reliable genetic overlap with creativity (**Supplementary Fig. 4** and **Supplementary Tables 13** and **14**). Interestingly, while SCZ did not demonstrate a significant genetic correlation with creativity in the LDSC analysis and SCZ PRS was only weakly associated with creativity, a considerable polygenic overlap (DC = 0.89) was suggested by the MiXeR analysis. Other psychiatric disorders such as MD, BD I, and AN, which showed weak genetic correlations with creativity in the LDSC analysis, also exhibited considerable polygenic overlap in the MiXeR analysis (DC = 0.94, 0.74, and 0.78, respectively). These results suggest that a substantial portion of genetic variants shared between creativity and these psychiatric disorders may have opposite effects. Although research on ASD, TS, and AN is limited and results are mixed for MD and ADHD, associations between creativity and various psychopathologies have previously been reported, especially between mood disorders and SCZ^4–9^. The current findings supplement these previous studies and offer new and deeper insight into genetic relationships between creativity and psychiatric disorders.

Based on polygenic overlap findings, additional and shared genetic variants between creativity and SCZ, MD, BD I, ADHD, and AN were identified via condFDR and conjFDR approaches. We identified 69 additional loci for creativity that were not detected in our initial GWAS using psychiatric disorders as conditional phenotypes. Moreover, using creativity as a conditional phenotype, additional loci were found for SCZ (number of SNPs [*n*] = 49), MD (*n* = 35), BD I (*n* = 25), ADHD (*n* = 18), and AN (*n* = 12) in comparison to their respective GWASs (**Fig. 5c** and **Supplementary Tables 15-19**). Our findings regarding the additional loci identified using creativity or psychiatric disorders as associated phenotypes suggest that there may be a common genetic basis between creativity and the five psychiatric disorders. With conjFDR analysis, shared genomic loci between creativity and SCZ (*n* = 50), MD (*n* = 80), BD I (*n* = 26), ADHD (*n* = 33), and AN (*n* = 21) were identified (**Supplementary Table 27**), highlighting the shared genetic structure between creativity and these five psychiatric disorders.

We found that using an ML-based continuous phenotype of creativity could identify more robust and significant results than using a dichotomous phenotype of creativity. GWASs using a continuous phenotype may have higher statistical power than those using a dichotomous phenotype^44^. As our results indicated, creativity, likewise other psychological traits, is a polygenic trait with a continuum of phenotypes rather than a dichotomous phenotype. It is also noteworthy that our ML-based GWAS found a strong correlation with the GWASs using traditional, dichotomous definitions of creativity (**Supplementary Figs. 10–13**).

Although this study provides an insight into the biological background of creativity and its related traits, it has limitations. First, we used the baseline occupation of the UKB participants to define their creativity. While using occupation to assess one’s creativity has been reliably used in previous studies^4–6, 8^, various other methods to measure creativity, such as divergent thinking tests, self-report questionnaires, and product-based assessments, are also available^1, 7^. Since creativity is a complex construct that encompasses aspects of cognition, personality, and external factors, other methods of measuring creativity could be utilized in the future to validate our findings. Second, creativity is a polygenic trait that is influenced not only by genes, but also the environment. As our study was focused on investigating the genetic basis of creativity, there was a limited exploration of environmental factors. Thus, our results should not be used to predict the creativity of individuals, but instead should be comprehended as additional evidence for the genetic basis of creativity. However, it is noteworthy that our results remained identical after adjustment for education years, one of the important environmental factors. Third, we only studied European individuals from the UKB cohort; thus, future studies involving more diverse ancestries are warranted. As culture can impact the development of creativity in a society, additional GWASs should attempt to replicate our findings with cohorts of various ancestries and identify additional genetic variants associated with creativity to provide greater insight into its biological underpinnings.

In summary, creativity is a polygenic trait that has a complex underlying genetic architecture. SCZ, MD, BD I, ADHD, and AN showed the most significant genetic associations with creativity across five analyses (LDSC, PRS, MiXeR, condFDR, and conjFDR). Although previous literature on the relationship between creativity and ADHD and AN is limited and conveys mixed results, our results indicate a significant genetic association of creativity with ADHD and AN. These comprehensive results suggest that it is necessary to analyze and consider the genetic architecture of psychiatric disorders from various perspectives. Our results are also clinically relevant, as psychiatric patients could be educated about this association between creativity and psychiatric disorders and use it to their advantage. Although patients who are creative may experience more severe symptoms of psychiatric disorders, they can be made aware of the beneficial elements of creativity. Activities utilizing their creativity (e.g., art therapy or book clubs) can provide benefits of rehabilitation and improve their quality of life. Additionally, the findings of overlapping biological mechanisms between creativity and psychiatric disorders can be useful for understanding traits that are related to both phenotypes. Lastly, the investigation of the biological underpinnings of creativity is also important for understanding genetic influences in everyday human behavior.

## Materials and Methods

### Study population

The UKB was constructed as a large prospective cohort study of approximately 500,000 individuals aged 40-69 years recruited from 2006 to 2010 across the UK. All participants provided electronically signed informed consent. The UKB was approved by the National Research Ethnic Committee (REC reference 11/NW/0382), and this secondary research was conducted in accordance with the principles of the Declaration of Helsinki and its later amendments. The details of the UKB project can be found elsewhere (https://www.ukbiobank.ac.uk/about-biobank-uk.)

### Genotyping and quality control

A total of 487,409 samples were genotyped using the Affymetrix UK BiLEVE Axiom or Affymetrix UKB Axiom arrays (Santa Clara, CA, USA), comprising more than 800,000 genetic variants. For imputation, phasing and imputation processes were centrally performed by the UKB using SHAPEIT3^45^ and IMPUTE2^46^, respectively, based on the combination reference panels of the 1000 Genomes Project Phase 3 and UK 10K. The variant-level quality control (QC) was applied for exclusion metrics, such as variants with a call rate < 95%, minor allele frequency (MAF) < 1 × 10^−4^, and Hardy–Weinberg equilibrium *P* < 1 × 10^−6^. After imputation, we performed a stringent QC using the PLINK 1.90 software^47^, applying three filters as follows: 1) call rate < 95% (missingness > 5%), 2) MAF < 0.5%, or 3) imputation quality scores (INFO) < 0.4. Genotypes with a posterior call probability < 0.90 were considered missing. A total of 9,575,249 SNPs met the QC criteria. Five sample-level QC exclusion criteria, including non-Europeans, samples with sex discordance between reported and genetically inferred sex, putative sex chromosome aneuploidy, no sex information, and participants who withdrew from the UKB were applied to the imputed data.

### Genome-wide association analysis

We performed a genome-wide association analysis using an ML-based method called REGENIE v2.2.4 for the creative probability^48^, using a ridge regression method with a leave-one-out cross-validation scheme to prevent overfitting. Based on a previous study^26^, birth year, squared birth year, cubic birth year, sex, the interaction of sex with birth year, squared birth year, and cubic birth year, batch, array, and ten principal components (PCs) of genetic ancestry were adjusted for in the association analysis. A genome-wide significant threshold of *P* < 5 × 10^−8^ was used to identify variants associated with creativity. A Manhattan plot was generated using a code obtained from Github (https://github.com/kbsssu/ManhattanGG). Regional plots were generated using LocusZoom v1.3.0 (http://locuszoom.sph.umich.edu/locuszoom)^49^.

Independent significant SNPs with *r*^2^ < 0.2 and *P* < 5 × 10^−8^ were identified through LD clumping in PLINK^47^. Among the 37 identified significant SNPs, we selected the most significant SNP per locus (within 1 mega-base pairs) as the lead SNP. The SNP annotation was performed using ANNOVAR^50^ and implemented in FUMA v1.3.7^51^.

### Gene mapping, functional annotation, and pathway analysis

The eQTL analysis and functional annotation were performed using FUMA^51^. eQTL analysis was performed using the GTEx (https://www.gtexportal.org/home/datasets) database v8^52^. An FDR < 0.05 was used to define significant eQTL associations. The gene-based analysis was carried out to find biological pathways based on the GO Consortium^53^ using MAGMA implemented in FUMA^51^.

### SNP-based heritability and cell type-specific analyses

LDSC v1.0.1^54^ was used to estimate the SNP-based heritability of creativity. The European LD scores of the 1000 Genomes Project v3 were obtained from GitHub (https://github.com/bulik/ldsc). The variants at the MHC region were excluded and common autosomal variants with an MAF > 1% in the European population were included. Using LDSC-SEG^55^, cell type-specific analyses were conducted to prioritize phenotype-associated tissues or cell types.

### Genetic correlation

The cross-trait genetic correlation (*r*_g_) of creativity probability with other phenotypes was estimated using LDSC^54^. We downloaded publicly available European GWAS summary statistics of 117 health-related phenotypes (**Supplementary Table 10**). GWAS summary statistics for nine psychiatric disorders were additionally used to find shared genetic backgrounds with creativity. The summary statistics were controlled for quality; their INFO was > 0.8 and MAF was > 0.5%. The FDR correction was used for multiple test correction (117 traits). For neuroimaging phenotype data, we used GWAS results for volumes of the region of interest (ROI) in the brain and GWAS results for DTI of ROI (**Supplementary Table 11**)^56, 57^. The details of the different data, including brain volume and DTI measure, are provided as follows: https://biobank.ctsu.ox.ac.uk/crystal/crystal/docs/brain_mri.pdf. We applied the FDR correction for ROI and each DTI scalar (FA, MD, AD, RD, and MO).

### Polygenic risk scoring analysis

We calculated the PRS for creativity based on the GWAS summary statistics of nine psychiatric disorders using PRSice-2 v2.3.3^58^ (**Supplementary Table 31**). Independent SNPs with *r*^2^ < 0.1 within 1 mega-base pairs were extracted based on LD clumping. The PRS of each psychiatric trait was calculated based on the pruned SNPs identified per trait. A total of 13 different clump *P*-value thresholds (5 × 10^−8^, 1 × 10^−7^, 1 × 10^−6^, 1 × 10^−5^, 1 × 10^−4^, 0.001, 0.05, 0.1, 0.2, 0.3, 0.4, 0.5, and 1) were tested to examine the association between the PRS and creativity across different SNP sets. The R^2^ value indicates the explained variance in the creativity of UKB individuals as a function of the PRS of each psychiatric disorder.

### Polygenic overlap

The shared polygenic overlaps between creativity and nine psychiatric disorders were quantified using MiXeR v1.2.0^59^ (http://github.com/precimed/mixer) (**Supplementary Tables 13** and **14**). The univariate analysis provides the number of causally associated SNPs with each trait (polygenicity), the average magnitude of additive genetic associations across causal variants (discoverability), and model fit criteria such as AIC and Bayesian Information Criterion (BIC) based on log-likelihood optimization of GWAS z-scores. Since MiXeR models additive genetic effects on two traits as a mixture of four bivariate Gaussian components (variants with no effect on both traits, variants with an effect on either trait, and variants with an effect on both traits), the bivariate analysis calculates a ratio of shared variants to the total number of variants (DC) and model fit (AIC and BIC). Conditional Q-Q plots were generated to depict the cross-phenotype polygenic enrichment between creativity and the nine psychiatric disorders analyzed.

### Conditional and conjunctional false discovery rate analysis

The condFDR analysis^60^ was employed (https://github.com/precimed/pleiofdr) to identify additional loci associated with psychiatric disorders that satisfies the model selection criteria (AIC) conditional on creativity in MiXeR and to find loci associated with creativity conditional on each psychiatric disorder. The SNP detection ability for a trait was improved based on substantial genetic association with a conditional trait. We selected the original GWAS of SCZ^18^, MD^19^, BD I^20^, ADHD^21, 22^, and AN^21, 22^, which had excluded UKB samples or included only UKB samples of less than 10% to minimize the inflation from sample overlap. Additional loci that were not detected in the original GWAS analysis were identified as well. We also applied the conjFDR analysis^60^ to identify shared genetic loci between psychiatric disorders and creativity. The maximum of the two condFDR values, which were calculated for every SNP, was taken as the conjFDR value between two traits^60^. Both condFDR and conjFDR approaches were applied by excluding SNPs within an intricate LD structure where the four regions are as follows: (MHC region, chromosome 6:25,119,106–33,854,733 base-pairs [bps]; 8p23.1, chromosome 8:7,200,000–12,500,000 bps; microtubule associated protein tau region, chromosome 17:40,000,000–47,000,000 bps; and apolipoprotein E region, and chromosome 19:44,909,039–45912,650 bps). The FUMA^51^ was used to define independent genetic loci with condFDR < 0.01 or conjFDR < 0.05, with the default settings. To identify additional loci related to psychiatric disorders in the condFDR analyses, we examined the variants within 1 mega-base pairs of the lead SNPs from the original GWAS results. Finally, independent genetic loci defined by both condFDR and conjFDR analyses were mapped to the genes in brain tissue via eQTL mapping, and tissue specificity was subsequently tested for the mapped genes considering differentially expressed genes (both up-regulated and down-regulated) based on the GTEx databases v8^52^ in FUMA^51^. The additionally identified loci for each trait were used in functional annotations for both psychiatric disorders and creativity using the condFDR approach. The default settings were applied for the remaining options in the functional annotation step, and the MHC region was excluded.

### Comparison with GWASs using narrow or broad definitions of creativity

We additionally performed GWASs on traditionally creative occupations (narrowly defined artistic professions) and broadly defined artistic or scientific professions (**Supplementary Table 2**), adjusting for the same covariates as the initial GWAS, using REGENIE v2.2.4^48^. Among 40 broadly defined artistic or scientific professions, seven professions (architects, draughtspersons, artists, authors/writers, actors/entertainers, dancers and choreographers, and musicians) were also categorized as narrowly defined artistic professions. The remaining 33 professions were therefore considered as creative proxies and excluded from the GWAS analysis for narrowly defined creative occupations. The education years were also included as a covariate in the initial association analysis for creative probability as well as the association models of dichotomous creative phenotypes to evaluate the effect of educational attainment on creativity. The genetic correlation between GWASs using continuous and dichotomous creative phenotypes was calculated using LDSC^54^. Furthermore, we extracted significant SNPs with *P* < 5 × 10^−8^ from the results of each GWAS to compare the direction of effect sizes. Based on the initial GWAS of ML-based creative probability, we estimated Spearman correlation coefficients and standard errors of the effect sizes obtained from GWASs of narrowly defined artistic professions, broadly defined artistic or scientific professions, and ML-based creative probability adjusted for education years. We also calculated the PRS for creativity using the GWAS of narrowly defined artistic professions based on the GWAS summary statistics of nine psychiatric disorders using PRSice-2 v2.3.3^58^ to compare the direction of their associations with the initial GWAS.

## Data availability

The GWAS summary statistics for creativity can be downloaded from the GWAS Catalog. The data including brain region phenotypes are available from the UKB (https://www.ukbiobank.ac.uk) upon project application. The GWAS summary statistics for genetic correlation analysis are available from several different sources such as GWAS ATLAS (https://atlas.ctglab.nl/traitDB), GWAS Catalog (https://www.ebi.ac.uk/gwas/studies/), Psychiatric Genomics Consortium (PGC) (https://www.med.unc.edu/pgc/download-results/), International Sleep Genetic Epidemiology Consortium (ISGEC) (https://www.kp4cd.org/dataset_downloads/sleep), and MEGASTROKE (http://www.megastroke.org/acknowledgments.html). The UKB summary statistics data from Ben Neale Group, CTGlab, and Lee Lab analyzed using Scalable and Accurate Implementation of Generalized mixed model (SAIGE) can be downloaded freely from http://www.nealelab.is/uk-biobank, https://ctg.cncr.nl/software/summary_statistics, and https://www.leelabsg.org/resources, respectively. The GWAS summary statistics for body mass index, education years, triglycerides, low-density lipoprotein cholesterol, and total cholesterol are available via previous studies (**Supplementary Table 10**).

## Acknowledgements

None

## Supplementary Materials

### 1. Supplementary Figures

**Supplementary Figure 1:** Creative probability distribution by job category.

**Supplementary Figure 2:** Regional association plots for GWAS of creativity.

**Supplementary Figure 3:** Quantile-quantile plots for GWAS of creativity (*n* = 241,736).

**Supplementary Figure 4:** Shared polygenicity underlying creativity and psychiatric disorders.

**Supplementary Figure 5:** Tissue specificity associated with mapped genes for additional loci from the conditional FDR results for psychiatric disorders (schizophrenia, major depression, bipolar I disorder, attention deficit/hyperactivity disorder, and anorexia nervosa) given creativity.

**Supplementary Figure 6:** Manhattan plots of −log_10_ scale of conditional FDR values for creativity given psychiatric disorders (schizophrenia, major depression, bipolar I disorder, attention deficit/hyperactivity disorder, and anorexia nervosa).

**Supplementary Figure 7:** Tissue specificity associated with mapped genes for additional loci from the conditional FDR results for creativity given psychiatric disorders (schizophrenia, major depression, bipolar I disorder, attention deficit/hyperactivity disorder, and anorexia nervosa).

**Supplementary Figure 8:** Manhattan plots of −log_10_ scale of conjunctional FDR values between psychiatric disorders (schizophrenia, major depression, bipolar I disorder, attention deficit/hyperactivity disorder, and anorexia nervosa) and creativity.

**Supplementary Figure 9:** Tissue specificity associated with genes mapped for loci from the conjunctional FDR results between psychiatric disorders (schizophrenia, major depression, bipolar I disorder, attention deficit/hyperactivity disorder, and anorexia nervosa) and creativity.

**Supplementary Figure 10:** Manhattan and quantile-quantile plots for GWAS of narrowly defined artistic professions (*n* = 219,722).

**Supplementary Figure 11:** Manhattan and quantile-quantile plots for GWAS of broadly defined artistic or scientific professions (*n* = 241,736).

**Supplementary Figure 12:** Manhattan and quantile-quantile plots for the GWASs with additional adjustment for education years (*n* = 241,736).

**Supplementary Figure 13:** Scatter plots comparing effect sizes from the GWASs based on the significant SNPs.

**Supplementary Figure 14:** Polygenic risk scores for psychiatric disorders associated with creativity using narrowly defined artistic professions.

### 2. Supplementary Tables

**Supplementary Table 1:** Participant demographic characteristics

**Supplementary Table 2:** Creative probability of each occupational group and narrowly and broadly defined creative professions

**Supplementary Table 3:** eQTL results for creativity

**Supplementary Table 4:** Gene set analysis

**Supplementary Table 5:** Enrichment for heritability partitioned based on 53 functional genomic annotations

**Supplementary Table 6:** Results from the multiple-tissue of gene expression analysis using LDSC-SEG

**Supplementary Table 7:** Results from the multiple-tissue analysis of chromatin data (validation) using LDSC-SEG

**Supplementary Table 8:** Results from the immune cell type of gene expression using LDSC-SEG

**Supplementary Table 9:** Results from the central nervous system (Cahoy) cell type of gene expression using LDSC-SEG

**Supplementary Table 10:** Genetic correlation between creativity and other traits (non-neuroimaging traits)

**Supplementary Table 11:** Genetic correlation between creativity and neuroimaging traits

**Supplementary Table 12:** Polygenic risk score for psychiatric disorders associated with creativity using ML-based creative probability

**Supplementary Table 13:** Univariate analysis (MiXeR)

**Supplementary Table 14:** Bivariate analysis between creativity and psychiatric phenotypes (MiXeR)

**Supplementary Table 15:** Independent genomic loci associated with schizophrenia given creativity at condFDR < 0.01

**Supplementary Table 16:** Independent genomic loci associated with major depression given creativity at condFDR < 0.01

**Supplementary Table 17:** Independent genomic loci associated with bipolar I disorder given creativity at condFDR < 0.01

**Supplementary Table 18:** Independent genomic loci associated with attention deficit/hyperactivity disorder given creativity at condFDR < 0.01

**Supplementary Table 19:** Independent genomic loci associated with anorexia nervosa given creativity at condFDR < 0.01

**Supplementary Table 20:** eQTL mapping of additional genomic loci from the condFDR results for schizophrenia conditional on creativity

**Supplementary Table 21:** eQTL mapping of additional genomic loci from the condFDR results for major depression conditional on creativity

**Supplementary Table 22:** eQTL mapping of additional genomic loci from the condFDR results for bipolar I disorder conditional on creativity

**Supplementary Table 23:** eQTL mapping of additional genomic loci from the condFDR results for attention deficit/hyperactivity disorder conditional on creativity

**Supplementary Table 24:** eQTL mapping of additional genomic loci from the condFDR results for anorexia nervosa conditional on creativity

**Supplementary Table 25:** Independent genomic loci associated with creativity given psychiatric disorders at condFDR < 0.01

**Supplementary Table 26:** eQTL mapping to of additional genomic loci from the condFDR results for creativity conditional on psychiatric disorders

**Supplementary Table 27:** Independent genomic loci jointly associated with creativity and psychiatric disorders at conjFDR < 0.05

**Supplementary Table 28:** eQTL mapping to genomic loci from the conjFDR results for creativity and psychiatric disorders

**Supplementary Table 29:** Polygenic risk score for psychiatric disorders associated with creativity using narrowly defined artistic professions

**Supplementary Table 30:** Relation matrix between creativity and psychiatric disorders

**Supplementary Table 31:** Summary of psychiatric disorder datasets

